# Early life environments shape adult cardiometabolic health during rapid lifestyle change

**DOI:** 10.1101/2025.06.06.658350

**Authors:** Rachel M. Petersen, Rob Tennyson, Tan Bee Ting A/P Tan Boon Huat, Kar Lye Tam, Marina M. Watowich, Kamal Solhaimi bin Fadzil, Colin Nicholas, Izandis bin Mohd Sayed, Kee Seong Ng, Yvonne A L Lim, Vivek V. Venkataraman, Ian J. Wallace, Thomas S. Kraft, Amanda J. Lea

## Abstract

Early life environments can have long-lasting impacts on health and fitness, but the evolutionary significance of these effects remains debated. Two major classes of explanations have been proposed: developmental constraints (DC) explanations posit that early life adversity limits optimal development, leading to long-term costs, while predictive adaptive response (PAR) explanations posit that organisms use early life cues to predict adult conditions, resulting in detriments when adult environments do not match expectations. We tested these hypotheses using anthropological and biomedical data for the Orang Asli—the Indigenous peoples of Peninsular Malaysia—who are undergoing a rapid but heterogenous transition from non-industrial, subsistence-based livelihoods to more industrialized, market-integrated conditions.

Using questionnaire data, we show that this shift creates natural variation in the degree of similarity between early life and adult environments. Using anthropometric and health data, we find that, more rural, subsistence-based early life environments are associated with shorter stature but better adult cardiometabolic health. Applying a quadratic regression framework, we find support for DC but not PAR in explaining adult cardiometabolic health, echoing findings and conclusions from other long-lived species. Overall, our results suggest that early life conditions can provide additive protection against common health issues associated with urban, industrialized lifestyle exposure.

## INTRODUCTION

Across mammals, early life environments shape later life health and fitness-related traits, a phenomenon known as “developmental plasticity” or “early life effects” [1–5]. In humans, early life environmental challenges (e.g., inadequate nutrition, psychosocial stress, or physical trauma) are associated with an increased lifelong risk of developing many of our most common health problems, such as cardiovascular disease, diabetes, hypertension, kidney disease, osteoporosis, osteoarthritis, respiratory disease, cancer, dementia, and depression [6–15]. Not surprisingly, early life adversity also predicts premature death in both human and non-human species [16,17] (but see [18]). Thus, across diverse organisms and populations, individuals who develop in favorable environments consistently do better later on, relative to conspecifics whose early lives were difficult in some way.

Two main classes of evolutionary hypotheses have been proposed to explain the correlation between early life environmental quality and adult outcomes. First, *the predictive adaptive response (PAR) hypothesis* posits that there has been selection for organisms to use environmental cues during development to alter their phenotype in ways that will be beneficial in a future, predicted adult environment that is similar [19,20]. Individuals that exhibit these “predictive adaptive responses” (also called “external PAR”, “adaptive developmental plasticity”, or “anticipatory maternal effects” [21,22]) are expected to exhibit better adult outcomes relative to individuals who experienced the same early life environment but made no phenotypic adjustments. Under this hypothesis, adult phenotypes should be optimal when conditions predicted from the early environment “match” the realized adult environment; in contrast, phenotypic “mismatch” and its associated costs will be more likely to occur when developmental cues do not accurately predict the adult environment.

In contrast to the PAR hypothesis, the *developmental constraints (DC) hypothesis* (also known as the “immediate adaptive response” or “silver spoon” hypothesis) emphasizes the immediate benefits but long-term costs of “making the best of a bad job” in adverse environments. This hypothesis posits that, when faced with suboptimal developmental conditions, organisms make somatic adjustments to ensure critical functions and immediate survival. However, these adjustments may come at the cost of compromised long-term health, fertility, or longevity [3,23]. Under this hypothesis, early life quality should thus predict adult outcomes regardless of the adult environment. We note that there are many derivatives of both the PAR [22,24–29] and DC [30–32] hypotheses, but we focus on PAR and DC because they are the most frequently discussed in the literature and because they are empirically testable. Importantly, while PAR and DC are sometimes discussed as alternative explanations, they are not mutually exclusive: for example, early life challenges could have long-term effects but be somewhat ameliorated in matched adult conditions [33–36].

The PAR hypothesis is especially relevant to cardiometabolic health in our species, because it grew out of observations in humans that early life nutritional stress was often associated with later life obesity and metabolic dysfunction [37]. This led to speculation that individuals who were undernourished in early life developed phenotypes that would have been well suited to scarcity in adulthood; when they instead found themselves in a plentiful adult environment “mismatched” to their expectations, health problems ensued. However, these results are also compatible with a DC interpretation, and simple negative associations between early life environmental quality and adult health are not adequate to disentangle the two models [20,38,39]. Instead, to test PAR and DC processes, it is necessary to examine cardiometabolic health-related traits in individuals from high- and low-quality early environments, when each of these sets of individuals experience high- and low-quality adult conditions [1–3,40–42]. Yet such datasets are rare, especially in humans [43–45], because early life and adult environments are often highly correlated and experimental manipulations are infeasible and unethical. When datasets containing the necessary diversity have been compiled in non-human animals and a handful of human populations (for cardiometabolic or other fitness-related traits), the results have generally favored DC over PAR interpretations (reviewed in [1,2,41]). However, more data are needed from humans specifically to test how, why, and under what conditions early life environments will alter later life cardiometabolic health. We also note that the statistical methods for testing DC and PAR models have improved recently [46], providing further motivation for additional studies.

To evaluate DC and PAR models, we focused on between- and within-individual environmental variation in a rapidly transitioning population—the Indigenous peoples of Peninsular Malaysia, known collectively as the Orang Asli. Traditionally, Orang Asli live in small, remote rainforest villages and subsist on some combination of hunting, gathering, fishing, and small-scale horticulture [47,48]. Over the last half-century, however, Malaysia has undergone one of the fastest rates of socioeconomic development in the world; this has caused many Orang Asli groups to experience varying types and degrees of lifestyle change including market integration (participation in the cash economy), acculturation (adoption of practices from the dominant culture), and urbanization (interaction with infrastructure associated with major towns or cities) [49–56]. At one extreme, some Orang Asli communities still remain isolated and continue to adhere to traditional lifestyles, but at the other extreme, some communities are now entirely bound by urban areas and fully integrated into the market economy. This has led to a situation in which 1) there is extreme variation in early life conditions within a single population and 2) some individuals experience similar early life and adult conditions, while others experience “mismatched” environments within their lifetimes.

As part of a long-term study of lifestyle change, health, and well-being among the Orang Asli [57], we collected in depth questionnaire data from 1155 adults on their early life and current environments, and we measured a suite of phenotypic traits in adulthood focused on cardiometabolic health (Supplementary Table 1). We used these integrative datasets combined with recently proposed methods for evaluating PAR and DC [46], with the overall goals of 1) understanding the patterns of rapid lifestyle shifts in this population, 2) testing for early life effects on later life outcomes, and 3) evaluating evolutionary hypotheses of early life effects.

## METHODS

### The Orang Asli of Peninsular Malaysia

The Indigenous Orang Asli comprise <1% of the population of Malaysia (∼238,000 people) and include at least 18 distinct ethnolinguistic groups, which are typically divided into three broad categories: the Semang/Negrito (traditionally nomadic hunter-gatherers speaking northern Aslian languages), Senoi (traditionally horticulturalists speaking central Aslian languages), and Proto-Malay (traditionally practitioners of mixed subsistence economies speaking Melanesian language dialects) [47,48]. Genetic variation exists between Orang Asli ethnolinguistic groups, especially the three major subgroups, but all are very similar genetically relative to surrounding Asian populations [58]. Over the last half-century, all ethnolinguistic groups have experienced different types and degrees of lifestyle change as a result of rapid socioeconomic development in Malaysia, with two trends having especially strong impacts.

First, expansion of industries focused on plantation agriculture (particularly oil palm and rubber) and natural resource extraction (particularly timber, tin, and petroleum) [59–61] have led to significant deforestation [62–66]; this has prompted an increased dependence on wage labor and the cash economy among many Orang Asli [49,50] as traditional lands are threatened.

Second, efforts by the Malaysian government to assimilate the Orang Asli into mainstream culture [48–50], including through development and resettlement schemes focused on “modern” facilities [49,50], have created shifts toward more urban, acculturated environments in certain geographic areas [51,53–55]. Importantly, due to the pace of these changes, variation exists in how long people have engaged in wage labor or lived in urban environments, with some people having made these transitions only recently and others having been exposed to such contexts for their entire lives. Further, some ethnolinguistic groups and individual villages have resisted sedentarization, interaction with the cash economy, and the authority of industry, government, and other entities more so than others [49,67]. Consequently, there are currently remote communities located in the rainforest that rely heavily on the availability of natural resources [68,69], as well as communities now immersed in fully industrialized economies and urban environments. Additional detail about the Orang Asli and lifestyle trends are provided in the Supplementary Materials.

### Questionnaire and cardiometabolic health biomarker data collection

The Orang Asli Health and Lifeways Project (OA HeLP) is an international research team of anthropologists, biologists, and physicians focused on documenting this lifestyle variation and its effect on health. A detailed protocol for OA HeLP is provided elsewhere [57], but data collection includes diverse information from questionnaires, health assessments, and biospecimens. The questionnaire, cardiometabolic health, and stature data used for this study were collected between June 2022 and December 2024 from Orang Asli individuals who were 18 years or older. During this time, researchers visited Orang Asli villages throughout Peninsular Malaysia. At each location, informed consent was collected at multiple levels: first by describing the project to the community as a whole and seeking the permission of community leaders, and subsequently through individual review of the protocol followed by formal, written consent. For this study, we worked with communities spanning the following ethnolinguistic groups: Batek, Jahai, Jakun, Kensiu, Mah Meri, Mendriq, Semai, Semaq Beri, Semelai, Temiar, and Temuan (Supplementary Table 2).

Data collection involved surveys about both early life and current environments, especially as these environments relate to lifestyle, acculturation, market-integration, and urbanization (a full list of questions is available in Supplementary Table 3). We also collected basic biomarkers of cardiometabolic health using the protocols described in [57] and in the Supplementary Materials, focusing on 7 body composition-related traits (body weight, body fat percentage, waist circumference, waist-to-hip ratio, body mass index (BMI), body roundness Index (BRI) [70], and obesity (Y/N)), 4 blood lipid traits (total cholesterol, HDL, LDL, triglycerides), and 3 blood pressure traits (systolic blood pressure, diastolic blood pressure, and hypertension (Y/N); see Supplementary Figure 1 and Supplementary Table 1 for trait-specific distributions and sample sizes). We included BRI in addition to the more standard BMI, because of mounting awareness that BMI can be a biased and misleading measure of adiposity [71]. To classify individuals as hypertensive (systolic BP >130 mmHg or diastolic BP >80 mmHg) and overweight or obese (BMI ≥25), we used standard cutoffs set by the American Heart Association. We note cutoffs for hypertension can vary globally [72], and we obtained very similar results when slightly higher cutoffs were used (systolic BP >140 mmHg or diastolic BP >90 mmHg [73]). For this study, we also analyzed 3 stature-related traits (standing height, hip height, and knee height), given previous work noting the sensitivity of these outcomes to the developmental environment [74–76]. Specifically, extensive prior research has noted the sensitivity of long bone and limb growth to early life conditions [74,75]. We note that we did not include stature-related traits in formal tests of the DC and PAR hypotheses, given a lack of theoretical motivation to do so.

### Quantifying early life and current environmental variation

To assess early life environments, we utilized data from retrospective surveys that covered multiple dimensions of lifestyle, urbanization, and acculturation, including dietary patterns, housing materials, parental subsistence activities, and household assets during childhood (Supplementary Table 3). Unordered, categorical answers were converted to binary variables noting a yes or no answer to each possible category (e.g., the question “when you were young, what was your father’s main work” was converted to a 0/1 variable for each possible type of work). Ordered, categorical answers were converted to a numeric scale (e.g., never, sometimes, often was converted to 1, 2, 3). Because some of our survey questions asked about livelihood strategies for an individual’s mother and father during childhood, we replaced missing values due to the death or absence of one parent with the values of the other parent. We implemented this procedure for 76 participants. After filling in this missing information about parental occupation, we also imputed data for 25 individuals that were missing answers to 1 or 2 out of the 30 remaining questions from the early life portion of the questionnaire. To do so, we used the R package “mice” and a Lasso approach [77]. This filled in and imputed dataset was then summarized via principal components analysis (PCA) using the “prcomp” function in R. We focused on the first component, which explained 21% of the variation. The decision to retain the first component was based on scree plot and biplot analyses to understand which individual variables loaded significantly on each PC: the first PC accounted for the largest and most interpretable portion of variation in early life environmental variables (see Supplementary Table 4 and Supplementary Figure 2).

We quantified current urbanicity in adulthood using a previously developed location-based score that we have validated in this population as a strong predictor of adult cardiometabolic health [78,79]. This score includes information about population density, access to infrastructure (e.g., electricity, sewage), household assets (e.g., televisions, mobile phones), subsistence activities, and education. These metrics are then averaged across individuals within a given village. This “urbanicity score” thus provides a relative measure of environmental exposure to urban infrastructure and market integration at the community level, which we have previously shown to be a stronger predictor of cardiometabolic health in this population than other possible indexes of acculturation, market-integration, and industrialization [78] (Supplementary Figure 3).

### Statistical analyses: exploring early life effects

To explore the relationship between early life environments and adult outcomes, we used linear models to examine associations between early life PC1 and continuous measures of stature and cardiometabolic health, and we used generalized linear models (binomial error distribution with logit link function) to assess relationships between early life PC1 and binary risk classifications (i.e., presence or absence of hypertension or obesity). In all models, we adjusted for current urbanicity score, age, and sex as covariates. Because the evolutionary models of interest suggest that early life effects on later life traits may depend on the current environment, we also included interactions between early life PC1 and current urbanicity score in all models. To account for multiple hypothesis testing, we applied a Benjamini Hochberg false discovery rate (FDR) correction [80] to each p-value set using the “p.adjust” function in R. We considered associations that pass a 10% FDR to be significant, but also note associations that pass a standard, nominal p-value of 0.05 in all analyses.

For all outcomes, we performed secondary analyses to consider two potential sources of heterogeneity in early life effects: ethnolinguistic group and sex. First, we used the “lme4” package in R [81] to run a set of models that included the major ethnolinguistic group categories as a random effect, to confirm that between-group variation in environment-outcome relationships did not impact our overall conclusions. These analyses included a four level factor for Senoi, Negrito, Proto Malay, as well as an “other” category covering mixed ancestry.

Second, we assessed the potential for sex-dependent effects of early life and current conditions by rerunning our main analyses after splitting the dataset by sex, as in [78]. For all analyses, we scaled our predictor and outcome variables in order to report standardized effect sizes.

### Statistical analyses: formal tests of DC and PAR

To formally test the predictions of DC and PAR models, focusing on our continuous cardiometabolic outcomes, we used a new regression-based approach proposed by Malani and colleagues [46]. This approach is based on the expectation that DC models would be supported by a positive relationship between early life environmental quality (*e*_0_) and adult health, while PAR models would be supported by a quadratic relationship between the difference in early life and adult environmental quality (*e_d_*) and adult health, where values closer to 0 (indicating “matched” early life and adult conditions) are associated with favorable health outcomes (Figure 1). Previous work has extensively validated this approach, showing it improves the sensitivity and specificity of inference of PAR and DC over other approaches [46]. Another benefit of this approach is that it allows both hypotheses to be tested simultaneously, and thus does not set them up as mutually exclusive.

**Figure 1.**
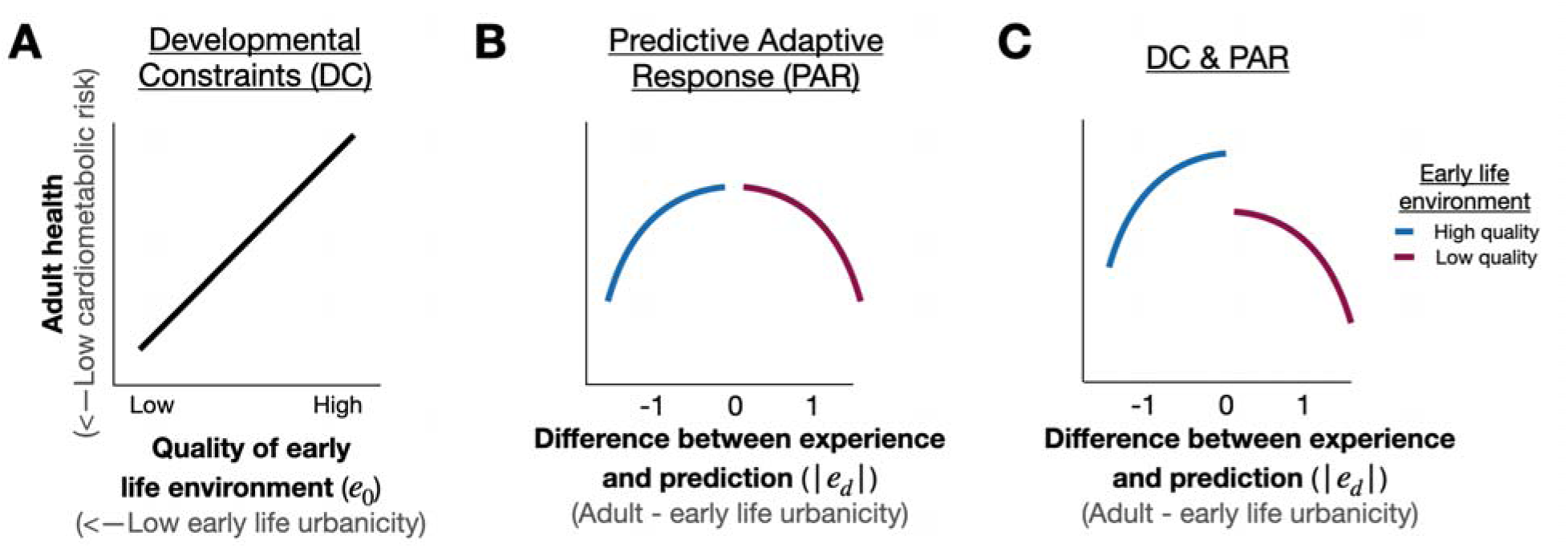
Conceptual framework for testing developmental constraints (DC) and predictive adaptive responses (PAR) using quadratic regression. (A) Hypothesized relationship between early-life environment and adult health under the DC framework, where low quality early-life conditions lead to worse adult health outcomes. (B) Hypothesized relationship between mismatches in early-life and adult environments and adult health under the PAR framework, where adult health is optimized when early-life and adult environments are matched. (C) Illustration of how both DC and PAR predictions can be simultaneously supported, with both low quality early-life conditions and mismatched environments leading to worse adult health outcomes.

Specifically, we used the following quadratic regression model as proposed by [46]:

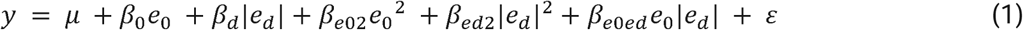

where *y* is a cardiometabolic outcome measured in adulthood, *e*_0_ is the developmental environment represented by early life PC1 and with effect size estimate *β*_0_, *e_d_* is the difference between the mean centered and scaled developmental and adult environments (with the adult environment represented by current urbanicity score). The absolute value of this variable is used, because we assume that the cost of erroneously predicting that the adult environment will be better than it really is, are approximately the same as the costs of erroneously predicting that the adult environment will be worse than it really is [46]. *β_d_* is the effect size estimate for the absolute difference between the early life and adult environments. Quadratic terms are included for both *e* and *e* (represented as *e*_0_^2^ and |*e_d_*|^2^, respectively) with their corresponding effect sizes (*β*_*e*02_ and *β*_*ed*2_), as is an interaction between *e*_0_ and *e_d_* (represented by *e*_0_|*e_d_*| with effect size estimate *β*_*e*0*ed*_ ), to allow for nonlinear relationships. *µ* is the intercept and *ε* represents residual variance not explained by the predictor variables. We note that all models also included covariates of age and sex, which are not represented in the above equation for simplicity.

To test for DC, we used the authors recommendation and provided code [46] to take the derivative of equation 1 for adult outcomes with respect to early life outcomes and test if this value is significant (via a Wald test) and positive. A positive derivative would indicate better early life conditions predict better adult health:

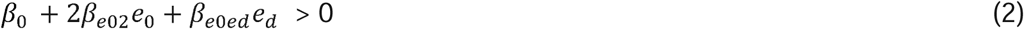

To test for PAR, we took the derivative of equation 1 of adult outcomes with respect to |e_d_ | and tested if this value was significant (via a Wald test) and negative. A negative derivative would indicate more similar early life and adult environments result in better adult health:

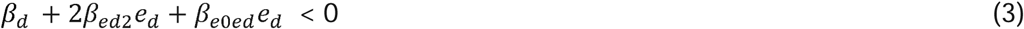

To perform these tests, we polarized outcome variables so that higher values represented “better” health (lower cardiometabolic disease risk) and predictor variables so that higher values represented hypothesized higher “quality” (i.e., more traditional) environments. We made this assumption based on previous work showing that more urban environments are consistently associated with worse cardiometabolic health [78]. Wald tests were performed using the “nlWaldTest” package in R. All p-values were corrected for multiple hypothesis testing using an FDR approach implemented with “p.adjust”.

All data processing, plotting, and statistical analyses were conducted in R version 4.3.

## RESULTS

### Extensive variation in early life environments in a transitioning population

We worked with 1155 Orang Asli individuals (738 females and 417 males, age range 18-91; Supplementary Figure 4) to characterize early life environments (Figure 2A) and their impacts on biomedical outcomes measured in adulthood (Supplementary Table 1), while controlling for their current environment (Figure 2B). As a result of past, ongoing, and heterogeneous development within Malaysia, we found extensive variation in early life conditions across space and time. For example, when analyzing the entire dataset, we found that only 20% of individuals born before 1975 lived in villages that were accessible by car or truck, but for individuals born after 2000, this percentage jumped to 45% (Figure 2C). This is strongly related to urbanization, as reliable roads coincide with the ability to transport goods and people both into and out of a location to interact with the market economy. Notably, there was substantial heterogeneity in early life infrastructure and the pace of change across villages: for example, in one urban location, 60% of older individuals compared to 100% of younger individuals grew up with road access, while in one peri-urban location, these numbers changed from 0% to 100% across generational categories, emphasizing rapid change within the last few decades (Figure 2D).

**Figure 2.**
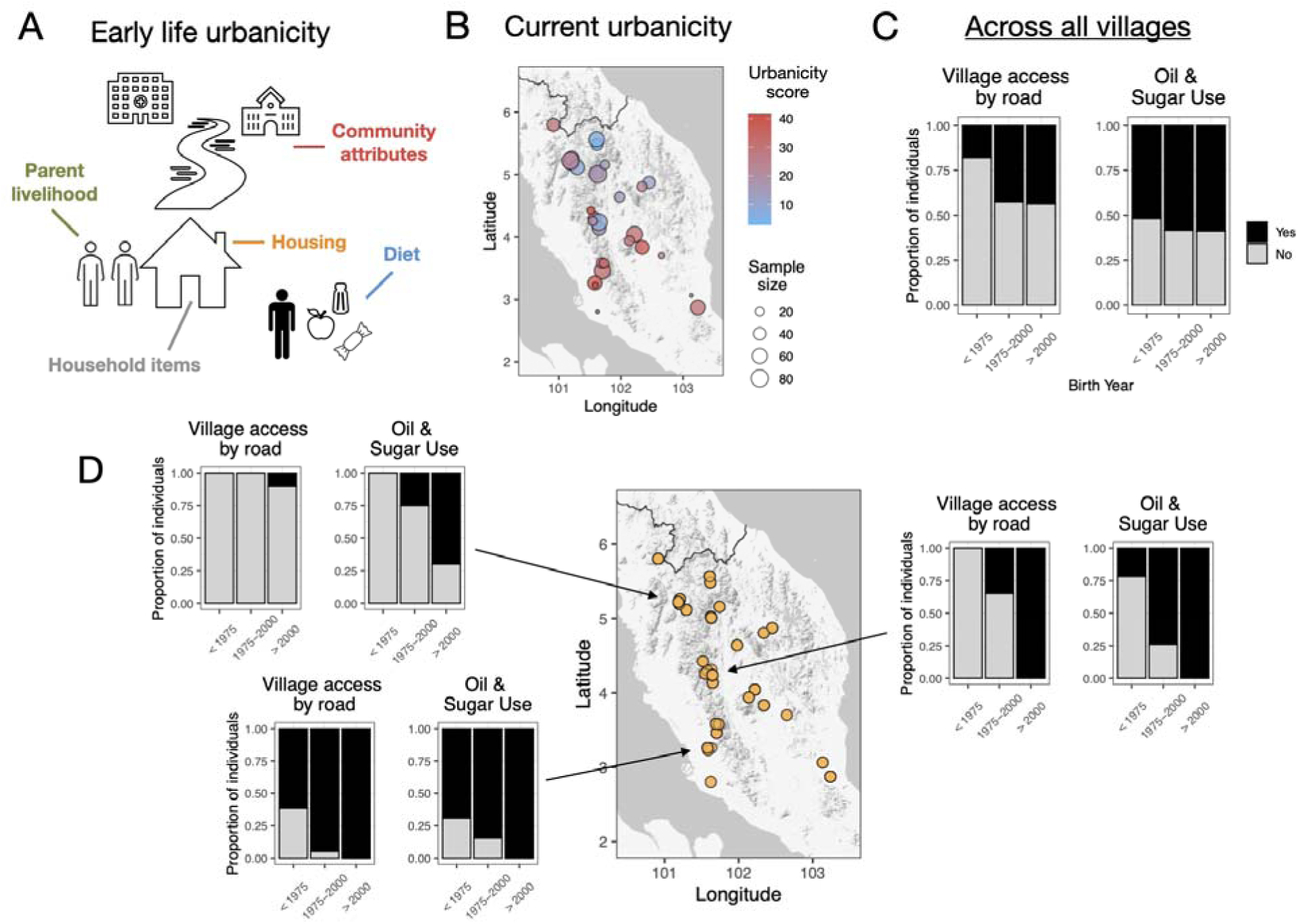
Study overview and variation in early life and current environments across Orang Asli villages. **(A)** Five major components of early life urbanicity quantified in our multidimensional retrospective survey. **(B)** Distribution of current urbanicity scores across participant villages (metric described in [78]). Greater values are associated with higher urbanicity and point size represents the number of individuals sampled at each locale. **(C)** Variation in two survey questions about early life environments, namely whether a participant’s childhood village was accessible by road to a car or truck and whether their household used oil and sugar. Results are summed across all surveyed villages and faceted by the individual’s birth year to show changes in the early life environment over time. **(D)** Between-village variation in how early life access to oil and sugar has changed over time, highlighting rapid and variable transitions from more traditional to more urban, market-integrated early life environments.

We summarized variation in early life environments using a principal components approach. We found that higher, positive values for PC1 primarily reflect access to market goods and urban infrastructure in early life, for example in the form of household possessions (e.g., ownership of a mobile phone, television), market-based foods (e.g., access to oil, sugar, or other foods from the store), and housing type (e.g., built of stone or wood materials). Whether an individual’s parents worked wage labor also loaded positively on PC1. In contrast, negative values for PC1 reflected more traditional possessions in early life, such as household ownership of instruments for subsistence activities (blowpipe, machete) and living in traditional bamboo housing. Thus, in general, we interpret higher scores to correspond to a more urban, more market-integrated, and less traditionally subsistence focused early life environment (Figure 3A).

**Figure 3.**
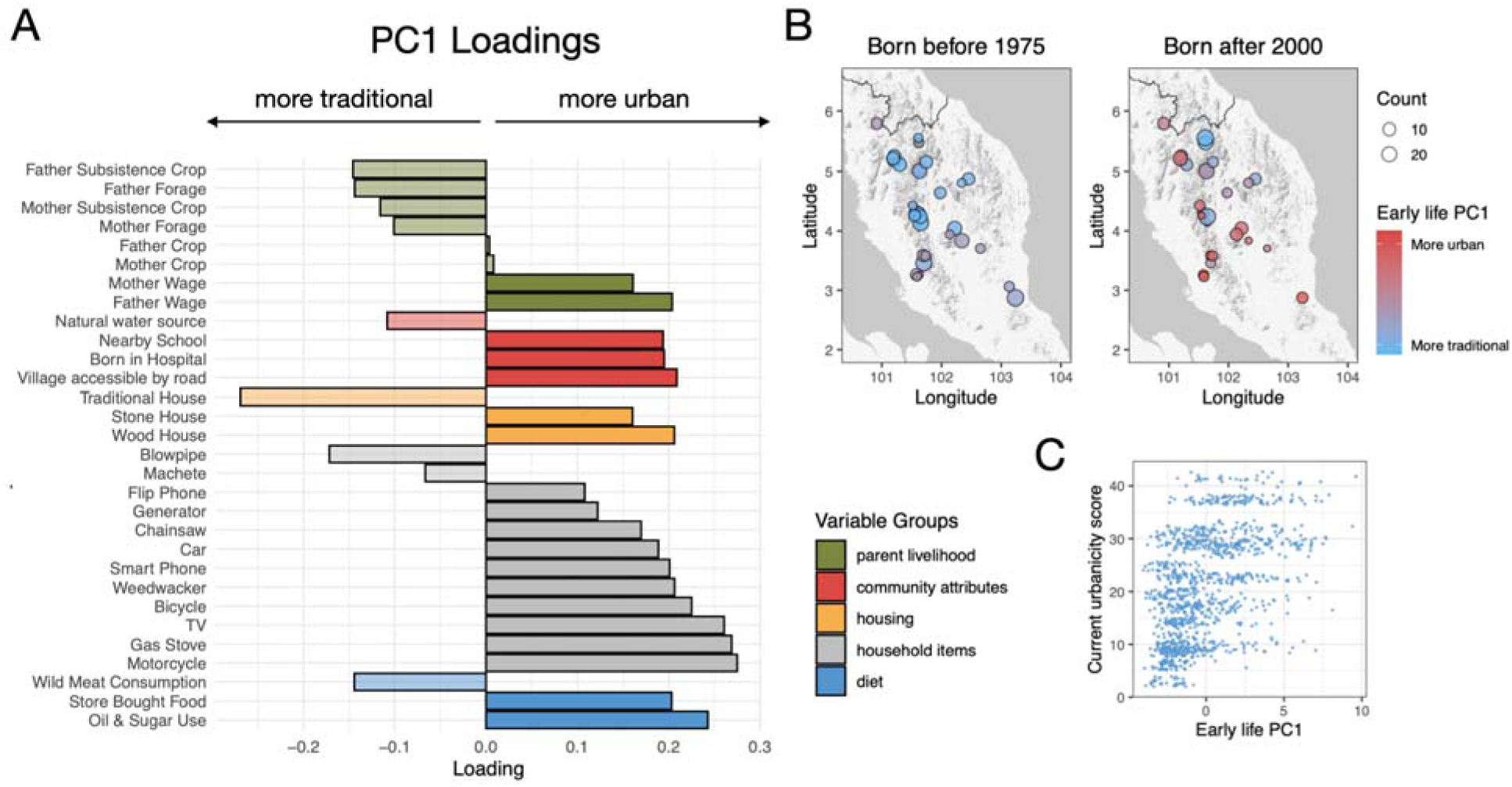
Early life urbanicity, market-integration, and industrialization summarized using a principal components approach. **(A)** Loadings of early life environmental variables on PC1 from a PCA used to derive the main axis of variation in early life environment. Higher loadings indicate stronger contributions to the early life urbanicity gradient, colors represent which of the five major components of early life environment each variable fell within, and transparency indicates whether the variable was expected to load negatively (lighter) or positively (darker) on PC1 based on a priori knowledge of the variable representing “traditional” versus “urban” conditions. **(B)** Distribution of early life PC1 scores across study villages for individuals born before 1975 (left) and born after 2000 (right), illustrating the rapid temporal change and spatial variation in early life exposures. **(C)** Comparison of individua early life PC1 versus current urbanicity scores, highlighting the noteworthy decoupling of these two factors in the Orang Asli, which facilitates testing the individual contribution of each to current health outcomes.

As expected for rapidly transitioning groups, we found that average early life conditions changed across time (Figure 3B), and that there were weak correlations between early life conditions and current urbanicity (linear model: R^2^=0.23, p<10^-10^; Figure 3C). This relationship to some degree reflects the fact that individuals who grew up in rural environments are now experiencing both rural and urban environments in adulthood, but individuals who grew up in urban environments are more likely to remain in urban environments. Further, early life-current environmental relationships exhibited clear age structure: older individuals were more likely to have grown up in early life environments that were more subsistence-focused and less market-integrated (linear model: R^2^=0.12, p<10^-10^) and correlations between early life PC1 and current urbanicity score were also somewhat age-dependent (linear model: R^2^=0.34, age x early life PC1 interaction term p=0.011; Supplementary Figure 5).

### Early life environments are associated with variation in cardiometabolic health and stature

Using models that controlled for age, current urbanicity score, sex, and interactions between early life and adult conditions, we found that early life conditions mattered for adult body composition, with exposure to more urban, market-integrated environments predicting higher body weights and body fat percentages, as well as a greater likelihood of meeting the criteria for obesity (FDR adjusted p-value<0.1); early life market-integration and urbanicity also trended toward predicting greater adult BMI (p<0.05; Figure 4A and Supplementary Table 5). When we examined stature-related traits, we found significant impacts of early life experience: growing up in a more urban, market-integrated environment was associated with greater standing height, hip height, and knee height (FDR adjusted p-value<0.1; Figure 4A and Supplementary Table 5). We further confirmed the effect of early life conditions on knee height in follow up models that included standing height as a covariate (p-value for early life PC1=0.027), to isolate proposed early life impacts on relative limb growth controlling for overall growth [82–84]. All reported results remained largely similar when controlling for potential heterogeneity between the three major ethnolinguistic groups (in these models, hip height, knee height, and body fat percentage passed a 10% FDR, while standing height and body weight passed a nominal p-value of 0.05; Supplementary Table 6). Sex-stratified models indicated that early life effects on stature may be stronger in males, while early life effects on body composition may be stronger in females; nevertheless, all effects were consistent in direction between the sexes (Supplementary Table 7 and Supplementary Figure 6)

**Figure 4.**
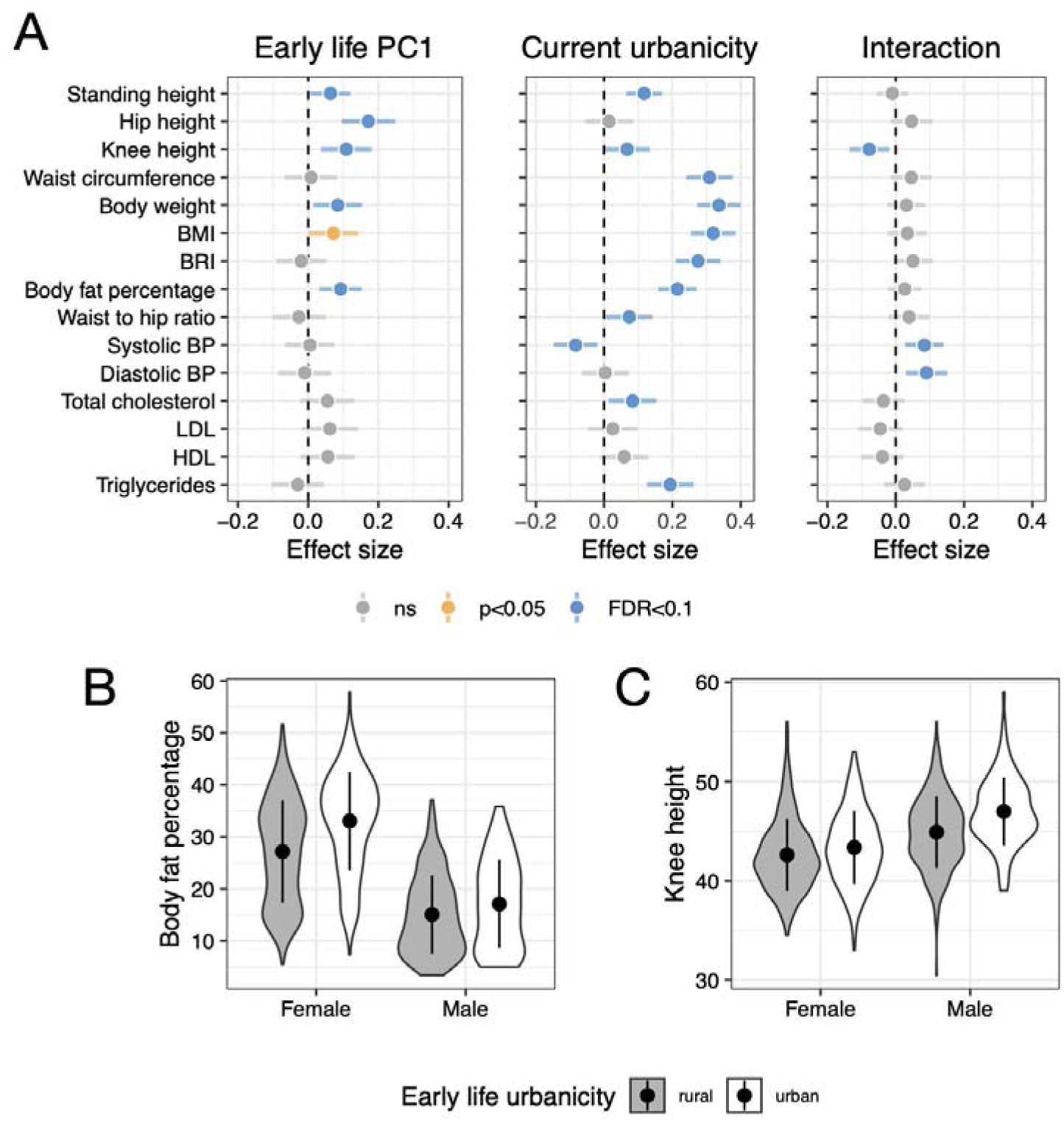
Effects of early life and current urbanicity on adult health outcomes. **(A)** Model results testing how early life urbanicity (left panel), current urbanicity (middle panel), or the interaction between the two (right panel) predicts health outcomes. Points represent model estimates, error bars represent 95% confidence intervals, and color indicates statistical significance (orange: p<0.05; blue: FDR-adjusted p-value<0.1). **(B-C)** Associations between early life urbanicity and example health outcomes. Individuals with more urban early life environments (white= PC1 > 0) exhibit significantly higher body fat percentage and greater knee height than those from more rural backgrounds (grey= PC1<0). Points indicate group means, error bars represent ±1 standard deviation, and violin plots show the distribution of the raw data uncorrected for other covariates.

We also identified strong effects of current urbanicity for 12/17 outcomes, consistent with our previous work [78]. Living in a more urban, market-integrated environment at the time of biomedical assessment was associated with greater adiposity (e.g., waist circumference, body weight, BMI), higher lipid levels (e.g., triglycerides, total cholesterol), and taller stature (e.g., standing height, knee height, FDR adjusted p-value<0.1; Figure 4A and Supplementary Table 5). The effects of the current environment were generally larger in magnitude than those of the early life environment: the one exception was knee height, which has previously been used as a biomarker of infancy and early childhood nutrition [85,86]. While interactions between early life and current conditions were generally positive, only 3 measurements passed our multiple hypothesis testing correction: knee height, systolic blood pressure, and diastolic blood pressure (FDR adjusted p-value<0.1; Figure 4A and Supplementary Table 5).

### Early life effects on adult cardiometabolic health are consistent with DC but not PAR

We used a recently proposed quadratic regression-based method to explicitly test for evidence of DC and/or PAR hypotheses [46] (Figure 1), focusing on cardiometabolic outcomes. With this modeling approach, we again found evidence that more urban, market-integrated early life conditions predicted worse adult cardiometabolic health, specifically larger waist circumference, as well as higher body weight, BMI, BRI, body fat percentage, total cholesterol, and triglycerides (Supplementary Table 8; FDR adjusted p-value<0.1 for the term). Explicit tests for DC—focused on asking whether there was a positive derivative between linear and non linear early life terms and adult outcomes—revealed similar conclusions and significant support for DC for the same body composition traits (waist circumference, body weight, BMI, BRI, body fat percentage) and lipid traits (total cholesterol, and triglycerides; Figure 5A&B and Supplementary Table 9). We found no evidence that experiencing more similar early life and adult conditions predicted better adult health (Supplementary Table 8; all p>0.05 for the term). Similarly, explicit tests for PAR—focused on asking whether there was a negative derivative between linear and non linear delta terms and adult outcomes—did not yield any significant results (Figure 5A&C and Supplementary Table 9).

**Figure 5.**
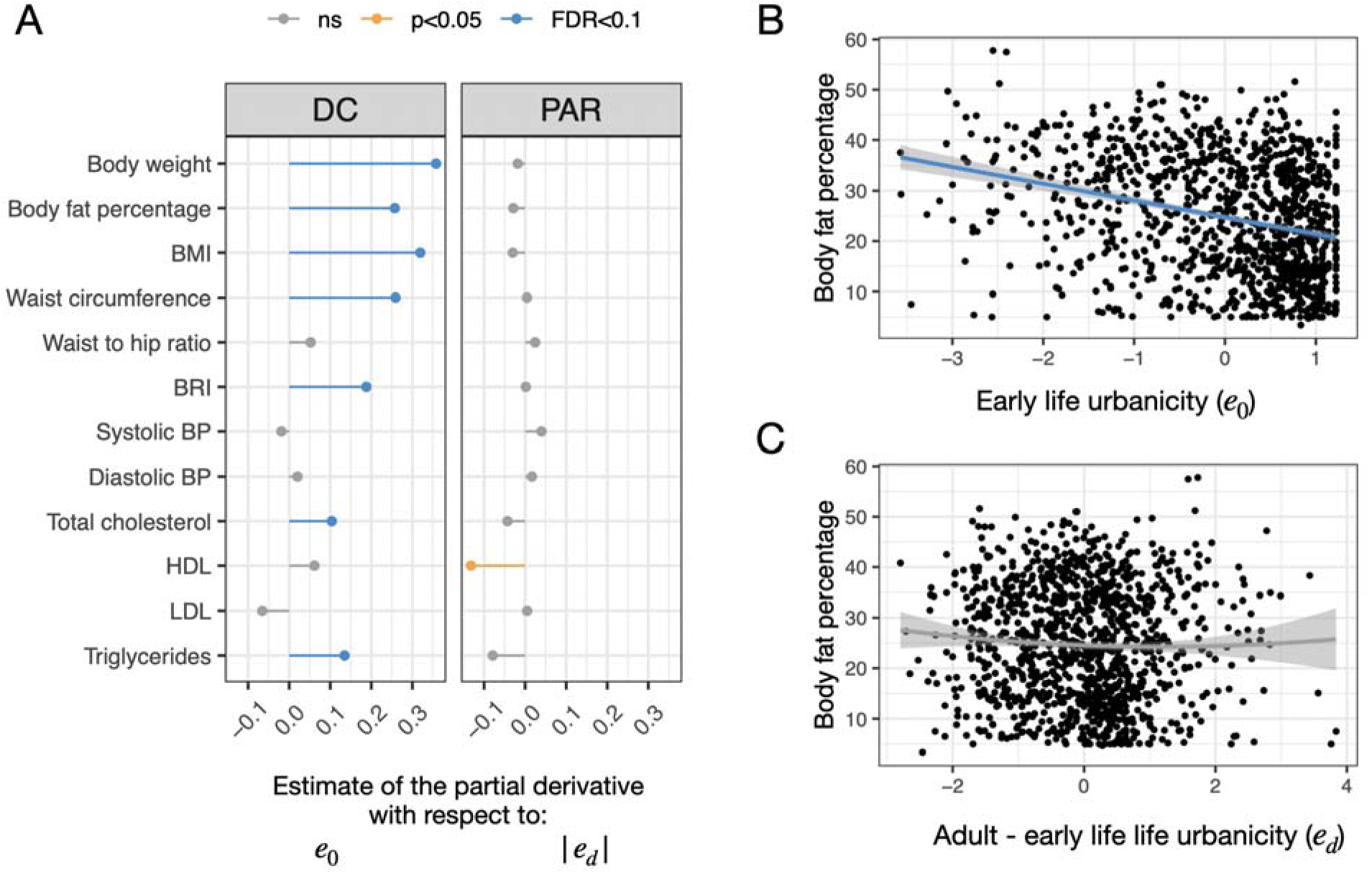
Results from quadratic regression tests for developmental constraints (DC) and predictive adaptive responses (PAR). Regression test results show support for the DC hypothesis on adult cardiometabolic traits. **(B)** As an example trait, we observe a strong linear relationship between early life environmental quality ( ) and adult body fat percentage, consistent with the predictions of the DC hypotheses, and **(C)** no evidence of a quadratic relationship between the difference in early life and adult environmental quality ( ) and adult body fat percentage, providing no support for the PAR hypothesis. Lines represent model predictions, shading represents the 95% confidence interval, and points represent the raw data.

## Discussion

Early life experiences can have profound and persistent effects on health and fitness-related traits in many mammalian species, including our own. Early life stressors consistently predict later life disease susceptibility and mortality rates, even if environmental conditions subsequently improve [1,2,11,17,87]. However, despite robust research linking early life environments to later life health outcomes in animal models and human cohorts in high-income countries [7,10,11], several gaps remain. Specifically, 1) considerably less research has focused on non-industrial contexts important to understanding human evolution [88] and 2) only a handful of studies have attempted robust tests of evolutionary explanations for early life effects in our species [43–45]. By focusing on a rapidly transitioning human population—the Orang Asli of Peninsular Malaysia—we found that variation in early life environments, ranging from more traditional, subsistence-focused lifestyles to more urban, market-integrated environments, has significant consequences: accounting for current environmental conditions, growing up in a more urban, market-integrated early life environment was associated with greater adult adiposity and greater adult stature. Our stature results are consistent with previous work in non-industrial groups: in the Indigenous Shuar as well as other populations living in the Southern Ecuadorian Amazon, early life market-integration was positively correlated with body size and growth in children and adolescents [75,89]. These results also echo work that has found positive effects of early life nutrition on linear growth and later life stature in other non-industrial groups [90,91], as well as relationships between early life resource availability and adult long bone length in wild non-human primates [92].

The environmental variation present within the Orang Asli also allowed us to test two evolutionary theories that have been proposed to explain variation in human cardiometabolic health [1–3]. Our formal tests of PAR and DC using recently proposed methods [46] provided clear support for DC: more urban, market-integrated early life environments predicted worse adult cardiometabolic health, regardless of the current environment. In contrast, we found no evidence that experiencing similar early life and current environments was associated with better outcomes in adulthood, as would be predicted by PAR. Practically, these results suggest that industrial transitions are unlikely to create health problems because of within-lifetime environmental mismatches; instead, our findings suggest that greater cumulative exposure to urban, industrialized environments across the life course are responsible for increasing rates of cardiometabolic disease [45]. We also note that our ability to create comprehensive measures of early life and current “urbanicity” represents a major advance over previous similar work [45], and allowed us to estimate continuous relationships between these variables and adult outcomes.

Our results add to empirical and theoretical studies that have generally favored DC over PAR interpretations in long-lived mammals [1], consistent with the simultaneously growing literature associating early life environmental adversity with compromised later life fertility, health, and mortality in these species (e.g., in bighorn sheep [93], spotted hyenas [94], wild baboons [95], and humans [10,96]). In theory, PAR should only be favored by natural selection when the environmental cue that triggers developmental plasticity is a reliable indicator of the adult environment, but this scenario is likely rare in long-lived organisms inhabiting dynamic environments [40,42,97]. In support, the best evidence of developmental plasticity consistent with PAR has been observed in short lived species, for example *Daphnia* [98], locusts [99], red squirrels [100], and voles [101,102]. Tests of PAR in long-lived species such as wild baboons [103], deer [104], and humans [43–45] have either rejected PAR explanations or identified weak evidence; additionally, some studies of long-lived species have found results that would not be consistent with either hypothesis (e.g., in African elephants [105], banded mongooses [106], wild baboons [33], and mountain gorillas [18]). Thus, while our results join previous studies in emphasizing that early life experiences often matter for cardiometabolic health in non-industrial and transitioning human populations [12,45,75,107,108], our evolutionary interpretation points toward these effects most often being additive rather than interactive with the current environment.

There are several limitations to our work. First, as with most non-experimental studies, early life and adult conditions, as well as their relationships with one another, are not randomly distributed across individuals. For example, early life PC1 and current urbanicity scores are weakly correlated (R^2^=0.23), driven by the fact that individuals born in urban environments are less likely to transition to more rural, subsistence-focused environments in adulthood than vice versa. We have done our best to control for known possible confounders (e.g., age), but note that there could be other unmeasured differences between individuals that occupy certain combinations of early life-current environments. Second, like many studies of early life effects in natural populations, the impacts of viability selection as well as differential adult mortality across environments are unknown. The role of viability selection has been previously noted as a challenge for tests of DC and PAR [104]. For example, individuals who experience early life challenges may be less likely to survive to adulthood, thus removing the most vulnerable individuals from the sample and creating a healthier surviving cohort. In our study, resource limitation, pathogen exposure, or reduced access to healthcare in rural areas could presumably lead to increased early life mortality in this group. In this scenario, our estimates of early life effects on later health would be conservative, likely underestimating the true differences that would exist in the absence of viability selection.

Our work also opens up several exciting new directions. First, developmental processes have received relatively little attention in studies of transitioning or non-industrial human populations [109]. Early life environmental components that have been well-studied in high-income countries—for example adverse childhood experiences [96]—may have different impacts across cultures and contexts, but this remains relatively unexplored. More work is thus needed to understand which, when, and how early life experiences shape adult outcomes in diverse human societies. This work could include more mechanistic studies, for example, of gene regulatory processes thought to embed early life experiences [110–112]. It could also include more detailed analyses of the timing and stability of early life effects across the life course. For example, in this study, we used relatively broad concepts of time (i.e., our survey instruments ask about conditions during childhood) as well as environment (i.e., summarizing dozens of questions with PCA), but future work could be more targeted.

Second, our results emphasize that early life exposure to market-integration, urbanization, and industrialization can have heterogeneous effects depending on the outcome, and this exposure is thus not unequivocally “bad”. For example, in this study, more urban early life environments predict “better” growth but “worse” cardiometabolic health. However, in work for example focused on the immune system, early life exposure to diverse microbes and pathogens is thought to potentially have some later life benefits (e.g., protecting against inflammation [108,113] or asthma [114]), but it can also carry costs [115]; further, this variable has complicated relationships with the rural-urban gradient [116]. Thus we emphasize that, depending on the outcome, it may be unclear what counts as a high versus low quality early life environment (when studying rural-urban gradients, or more generally). As a result, certain outcomes may not easily fit within the DC or PAR conceptual models. Moving forward, we are excited for studies that improve our understanding of the types of traits that have evolved to be sensitive to developmental conditions in our species, how tradeoffs between biological systems are induced by early life challenges [117,118], and ways we can use this knowledge to improve human health during lifestyle transitions [109].

### Ethics

Procedures for this study have been reviewed and approved by the Medical Review and Ethics Committee of the Malaysian Ministry of Health (protocol ID: NMRR-20-2214-55565), the Malaysian Ministry of Economy (permit ID: EPU 40/200/19/3911), the Malaysian Department of Orang Asli Development (permit ID: JAKOA.PP.30.052 JLD 21 (98)) and the Institutional Review Boards of the University of New Mexico (protocol ID: 14420) and Vanderbilt University (protocol ID: 212175). Throughout the project, we have followed established principles for ethical biomedical research among Indigenous communities, including fostering collaboration, building cultural competency, being transparent about research practices, supporting capacity building, and disseminating research findings [119].

## Data accessibility

OA HeLP’s highest priority is the minimization of risk to study participants. OA HeLP adheres to the ‘CARE Principles for Indigenous Data Governance’ (Collective Benefit, Authority to Control, Responsibility, and Ethics). OA HeLP is also committed to the ‘FAIR Guiding Principles for scientific data management and stewardship’ (Findable, Accessible, Interoperable, Reusable). To adhere to these principles while minimizing risks, individual-level data are stored in the OA HeLP protected data repository, and are available through restricted access.

Requests for de-identified, individual-level data should take the form of an application that details the exact uses of the data and the research questions to be addressed, procedures that will be employed for data security and individual privacy, potential benefits to the study communities and procedures for assessing and minimizing stigmatizing interpretations of the research results. Requests for de-identified, individual-level data will require institutional IRB approval (even if exempt). OA HeLP is committed to open science and the project leadership is available to assist interested investigators in preparing data access requests (see orangaslihealth.org for further details and contact information). Code to generate the current urbanicity score is available on GitHub (https://github.com/mwatowich/Multi-population_lifestyle_scales). All code for analyses presented here are also available on Github (https://github.com/rachpetersen/Early_life_effects_health_OA).

## Competing interests

The authors declare no competing interests.

## Funding

This work was supported by the National Science Foundation (BCS-2142090), the Canadian Institute for Advanced Research (Azrieli Global Scholars Program), and the Pew Charitable Trusts (Pew Biomedical Scholars Program). The funders had no role in study design, data collection and analysis, decision to publish, or preparation of the manuscript.

## Acknowledgements

We thank Orang Asli participants and communities for their interest, support, hospitality and willingness to work with us on this project. We thank the OA HeLP field team and volunteers, whose expertise and commitment made this work possible. We are grateful to the local organizations and institutions that we work with, namely the Center for Orang Asli Concerns, the Malaysian Red Crescent Society, Federation of Private Medical Practitioners’ Associations of Malaysia (Drs4All program), Hospital Orang Asli Gombak, and Universiti Malaya.

## Notes

### Competing Interest Statement

The authors have declared no competing interest.

