## Supplemental Methods and Figures for "Early life environments shape adult cardiometabolic health during rapid lifestyle change"

Supplemental materials for **“Early life environments shape adult cardiometabolic health during rapid lifestyle change”**

This supplement consists of:

**Supplementary Methods**

**Supplementary Figures 1-6**

- Supplementary Figure 1. Distributions for all measured outcomes.
- Supplementary Figure 2. Scree plot from a principal components analysis of early life environmental variation.
- Supplementary Figure 3. Distribution of the “urbanicity score” used to characterize the current environment.
- Supplementary Figure 4. Distribution of participant ages.
- Supplementary Figure 5. Rapid lifestyle transition creates relationships between age and environmental experiences.
- Supplementary Figure 6. Early life PC1 effects estimated from sex-stratified analyses.

**Supplementary References**

**Supplementary Methods**

The Orang Asli of Peninsular Malaysia

The population of Peninsular Malaysia is multiethnic, with an ethnically Malay majority and large minorities of Malaysian Chinese and Indians. The Indigenous Orang Asli comprise <1% of the population (~150,000 people) and include at least 19 distinct ethnolinguistic groups, which are typically divided into three broad categories: the Semang or Negrito (traditionally nomadic hunter-gatherers speaking northern Aslian languages), Senoi (traditionally horticulturalists speaking central Aslian languages), and Proto-Malay (traditionally practitioners of mixed subsistence economies speaking Melanesian language dialects) [[1,2]](https://paperpile.com/c/Nzz26N/ZQK4E+oR3lt). Today, many Orang Asli people are fluent in both their native language and the lingua franca of Malaysia (Malay). Genetic variation exists between Orang Asli ethnolinguistic groups, but all are similar genetically relative to surrounding Asian populations [[3]](https://paperpile.com/c/Nzz26N/HaI8i).

Over the last half-century, Malaysia has undergone one of the fastest rates of socioeconomic development in the world, which has precipitated lifestyle changes among Orang Asli in numerous and complex ways. Two trends have had especially profound impacts. First, a key component of efforts to accelerate growth of the national market economy has been the expansion of industries focused on plantation agriculture (particularly oil palm and rubber) and natural resource extraction (particularly timber, tin, and petroleum) [[4–6]](https://paperpile.com/c/Nzz26N/RNg6C+EzHVI+BCfAW). As a result, Malaysia has experienced marked deforestation [[7–9]](https://paperpile.com/c/Nzz26N/hSkJS+A2gek+tIjkI), which has fragmented and destroyed a large fraction of the lands traditionally occupied by Orang Asli [[10–14]](https://paperpile.com/c/Nzz26N/PMkrj+pPU85+YErm5+wkpqI+zhSG9). Many Orang Asli have thus shifted their livelihoods to a dependence on wage labor, which has been associated with increases in market integration, acculturation, and urbanization [[15,16]](https://paperpile.com/c/Nzz26N/p5fkp+aTlNb). Second, a longstanding objective of the Malaysian government has been to promote the assimilation of Orang Asli into mainstream Malaysian society and the integration of Orang Asli economies with the national market economy [[2,15,16]](https://paperpile.com/c/Nzz26N/oR3lt+p5fkp+aTlNb). To this end, government programs have been established to resettle and regroup Orang Asli people into consolidated villages with “modern” facilities including administrative centers, schools, shops, clinics, prefabricated houses, and roadways [[15,16]](https://paperpile.com/c/Nzz26N/p5fkp+aTlNb). Village members are typically provided some form of income-generating activity (e.g., rubber trees). Ultimately, due to these two trends, many Orang Asli people currently live in highly acculturated, market-integrated contexts [[17–20]](https://paperpile.com/c/Nzz26N/DHMo9+RlI40+0741c+fTPTW). Yet, variation exists in how long people have lived in such contexts, with some people having transitioned only recently and others having lived in such contexts for their entire lives.

Notwithstanding the strength of industry, government, and other forces, many Orang Asli continue to adhere strongly to traditional lifestyles, albeit on lands that are threatened or degraded relative to those occupied by earlier generations. The most averse to change have been Negrito groups (e.g., Batek, Jahai), which traditionally practice a hunter-gatherer lifestyle [[15,21–23]](https://paperpile.com/c/Nzz26N/p5fkp+uPqe5+zdSOF+sW66w). Because of the traditionally nomadic, egalitarian, and autonomous nature of these groups [[21,24,25]](https://paperpile.com/c/Nzz26N/2DyDR+KBa2L+uPqe5), many have resisted sedentarization and the authority of industry, government, and other entities [[15,21]](https://paperpile.com/c/Nzz26N/uPqe5+p5fkp). Consequently, there still exist remote communities located in the rainforest that rely heavily on the availability of natural resources, including several in which members of our team have established strong relationships [[26,27]](https://paperpile.com/c/Nzz26N/PwALy+PY4Dc).

In addition, within just the last two decades, a distinct type of lifestyle change has been occurring among certain Orang Asli groups—a reversion from a highly acculturated, market-integrated lifestyle to a more traditional lifestyle. This trend has been driven primarily by dissatisfaction with various aspects of life in government-run resettlement/regroupment towns, and has been restricted to remote regions where transitioning to a more traditional, rainforest-based lifestyle is still possible. Although currently undocumented in the academic literature, this trend is well-known to researchers and agencies working with Orang Asli. For example, members of our team have established relationships with people from multiple Senoi (e.g., Semai, Temiar) rainforest communities, in which many inhabitants grew up in resettlement/regroupment towns but today subsist primarily on horticulture, hunting, and gathering. In sum, the Orang Asli represent a highly unique and ideal study system for this project, with individuals having experienced a range of different lifestyle trajectories.

The Orang Asli Health and Lifeways Project and data collection

A detailed protocol for the Orang Asli Health and Lifeways Project is provided in[[28]](https://paperpile.com/c/Nzz26N/fvI23). The interview and health data summarized here were collected from self-reported Orang Asli individuals who were 18 years or older. To do so, researchers visited locations throughout Peninsular Malaysia where Orang Asli individuals were known to reside. At each sampling location, the headman and community were first consulted about the project before individuals were invited to participate. After this, individual consent was obtained from each individual in their language of choice. Structured interviews were conducted with all participants to collect information about both early life and current experiences, especially as these experiences relate to lifestyle, acculturation, market-integration, and urbanization.

Participant height was measured with a stadiometer. Body weight and body fat percentage were measured with an electronic scale capable of gauging bioelectrical impedance (TANITA's BC-558 FDA Cleared Ironman Segmental Body Composition Monitor). Waist circumference, hip circumference, leg height, and knee height were measured with a tape measure. Blood pressure was measured with an Omron 3 Series Upper Arm Blood Pressure Monitor. Prior to these measurements, participants were resting in a seated position for at least 5 minutes, and we used the second of two measurements to account for a “white coat” effect. Blood lipids were measured from a venous blood draw using a CardioCheck Plus Analyzer. BRI was calculated as 364.2 − 365.5 × √(1 − [waist circumference in centimeters / 2π]2 / [0.5 × height in centimeters]2), according to the formula developed by [[29]](https://paperpile.com/c/Nzz26N/26sT). BMI was calculated as weight (kg)/height(m)^2^.

**Supplementary Figures**

**Supplementary Figure 1. Distributions for all measured outcomes.** Units for each trait are provided in Supplementary Table 1.

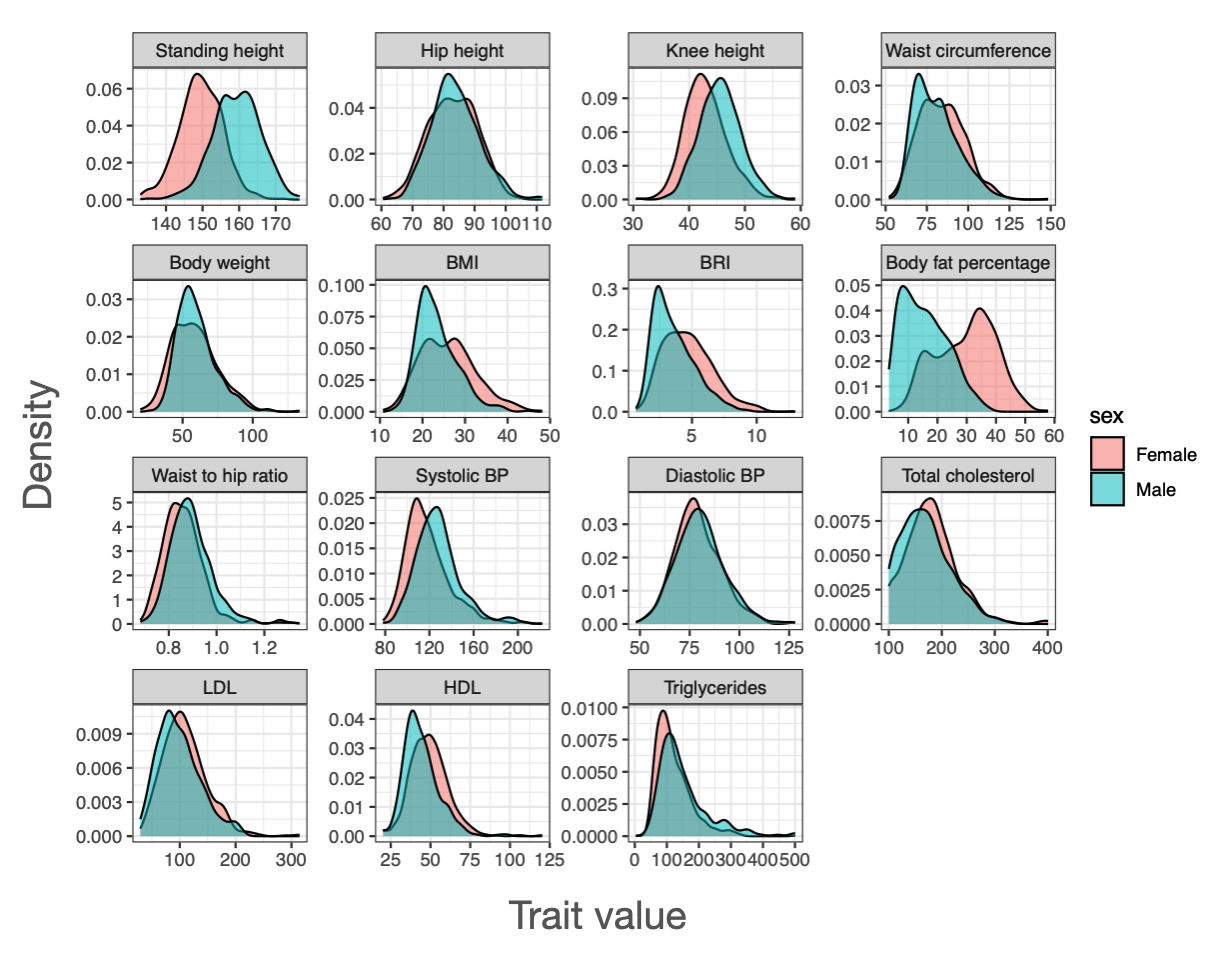

**Supplementary Figure 2. Scree plot from a principal components analysis of early life environmental variation.** Plot shows the top 10 principal components versus the proportion of variance explained by each component.

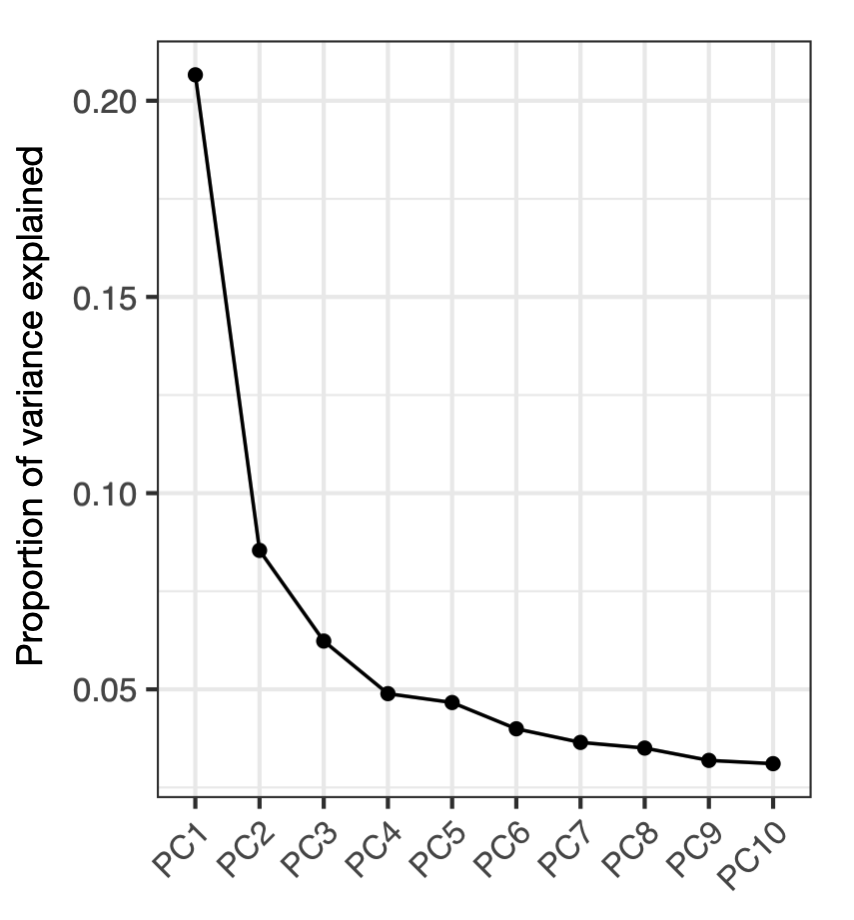

**Supplementary Figure 3. Distribution of the “urbanicity score” used to characterize the current environment.** The urbanicity score is described in [[30]](https://paperpile.com/c/Nzz26N/Z4x8).

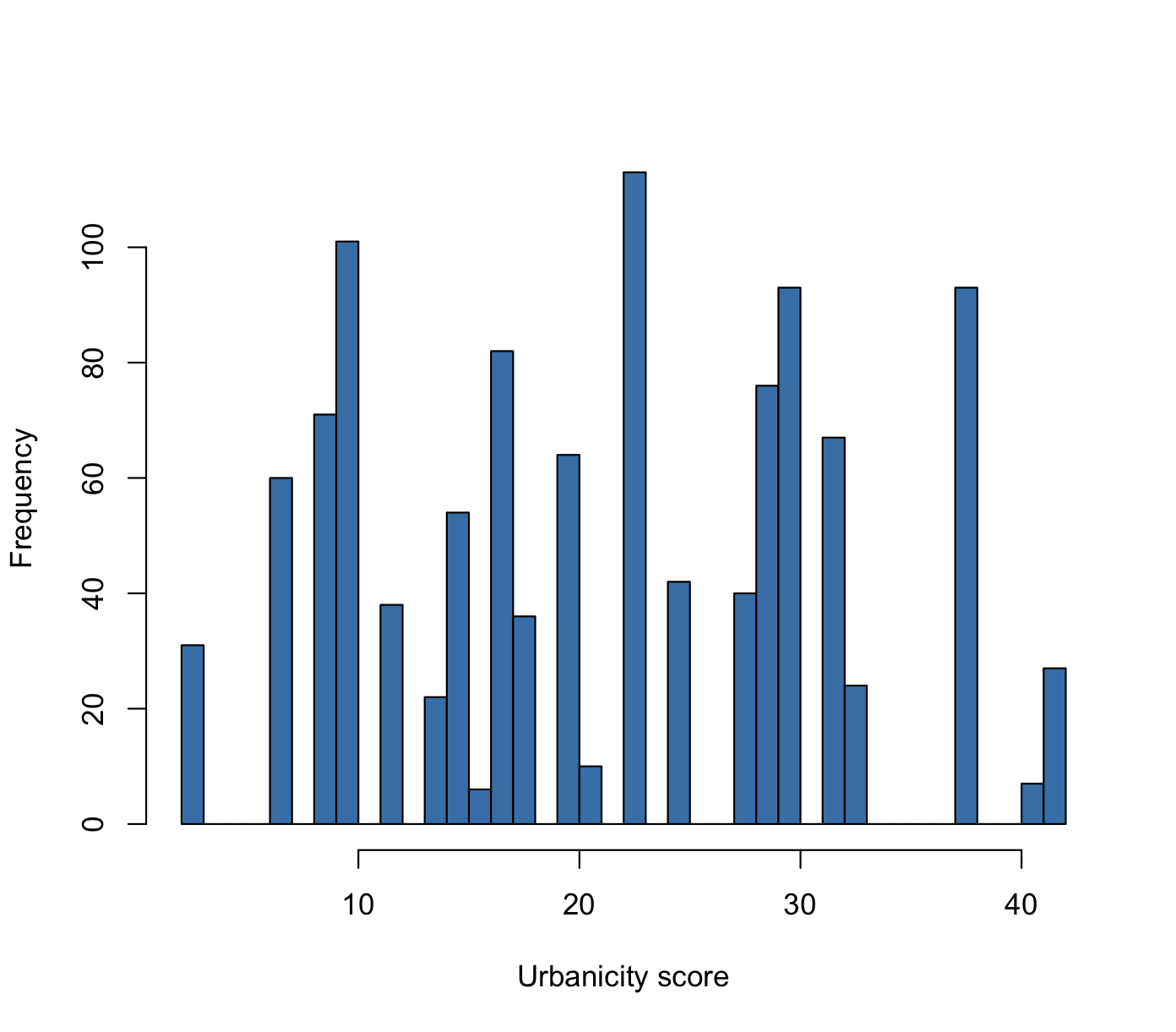

**Supplementary Figure 4. Distribution of participant ages.**

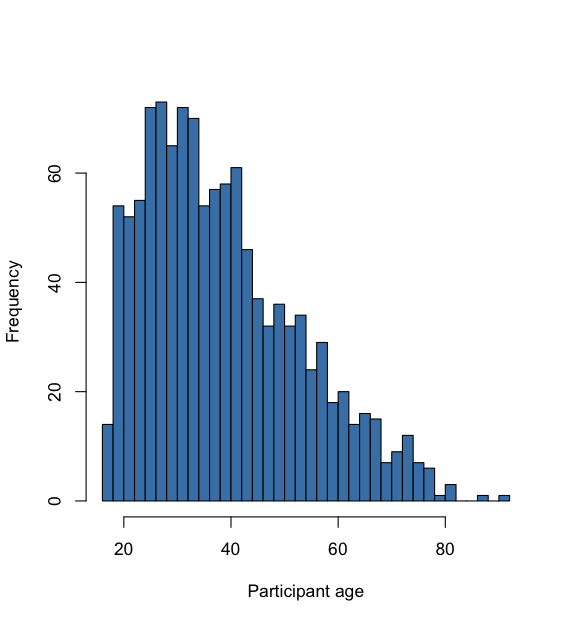

**Supplementary Figure 5. Rapid lifestyle transition creates relationships between age and environmental experiences.** A) Relationship between age and and early life PC1. Older individuals were more likely to have grown up in early life environments that were more subsistence-focused and less market-integrated (linear model: R^2^=0.12, p<10^-10^). B) Relationship between early life PC1 and current urbanicity, stratified by participant age group. The four age groups represent quartiles of the overall dataset. Relationships between early life PC1 and current urbanicity are somewhat age-dependent, though these dependencies appear heterogenous and not trending in any particular direction (linear model: R^2^=0.34, age x early life PC1 interaction term p=0.011).

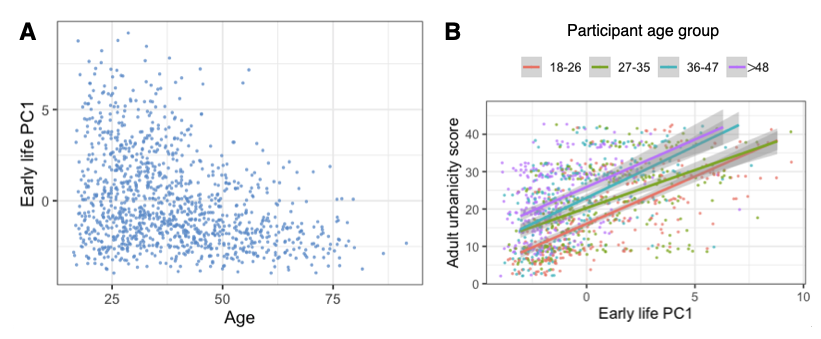

**Supplementary Figure 6. Early life PC1 effects estimated from sex-stratified analyses.** All models controlled for age and current urbanicity. Bars represent 95% confidence intervals. Orange bars represent effects that passed a nominal p-value threshold of 0.05, while blue bars represent effects that passed a 10% false discovery rate threshold.

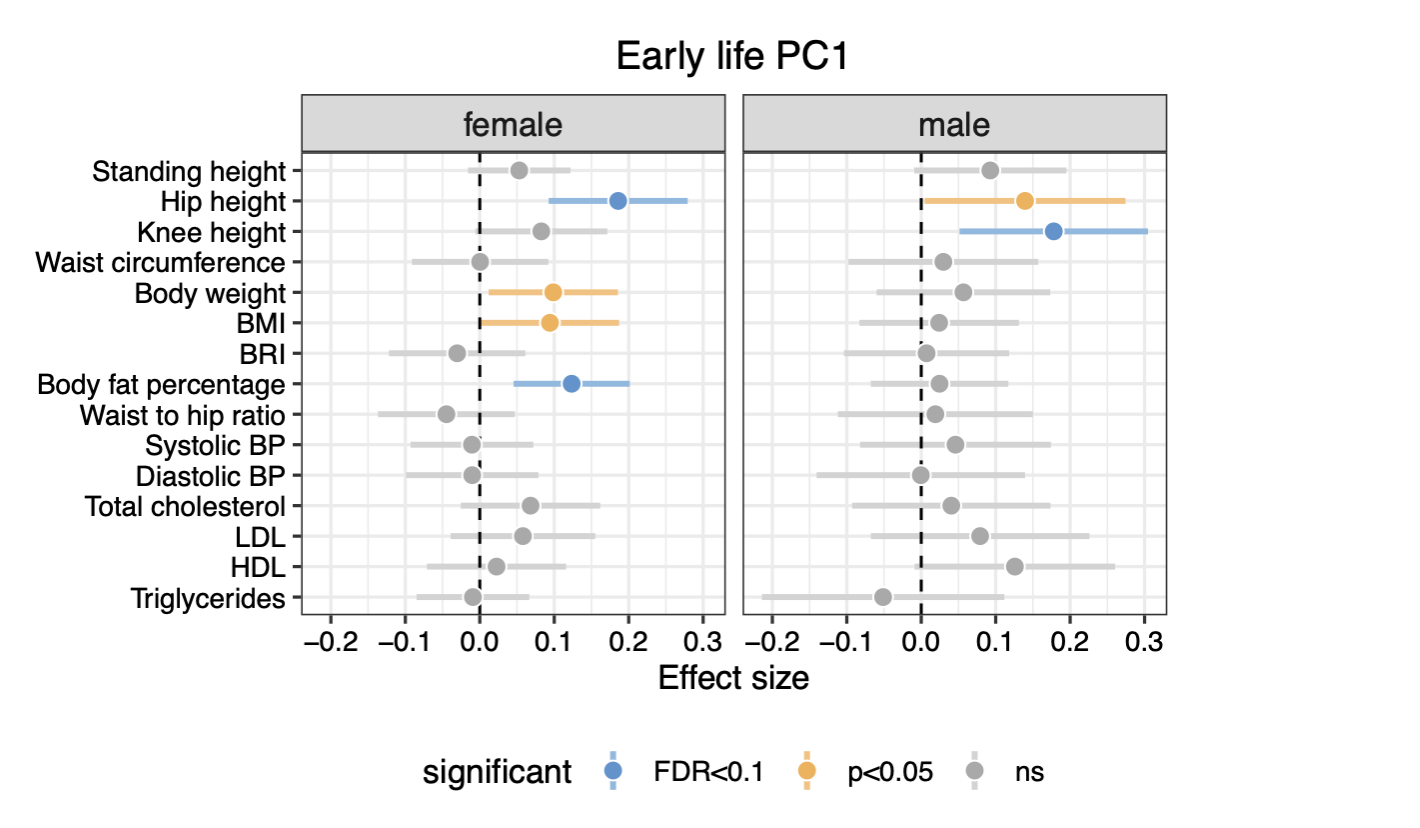
